# A descriptive marker gene approach to single-cell pseudotime inference

**DOI:** 10.1101/060442

**Authors:** Kieran R Campbell, Christopher Yau

## Abstract

Pseudotime estimation from single-cell gene expression allows the recovery of temporal information from otherwise static profiles of individual cells. This pseudotemporal information can be used to characterise transient events in temporally evolving biological systems. Conventional algorithms typically emphasise an unsupervised transcriptome-wide approach and use retrospective analysis to evaluate the behaviour of individual genes. Here we introduce an orthogonal approach termed “Ouija” that learns pseudotimes from a small set of marker genes that might ordinarily be used to retrospectively confirm the accuracy of unsupervised pseudotime algorithms. Crucially, we model these genes in terms of switch-like or transient behaviour along the trajectory, allowing us to understand why the pseudotimes have been inferred and learn informative parameters about the behaviour of each gene. Since each gene is associated with a switch or peak time the genes are effectively ordered along with the cells, allowing each part of the trajectory to be understood in terms of the behaviour of certain genes. In the following we introduce our model and demonstrate that in many instances a small panel of marker genes can recover pseudotimes that are consistent with those obtained using the entire transcriptome. Furthermore, we show that our method can detect differences in the regulation timings between two genes and identify “metastable” states - discrete cell types along the continuous trajectories - that recapitulate known cell types. Ouija therefore provides a powerful complimentary approach to existing whole transcriptome based pseudotime estimation methods. An open source implementation is available at http://www.github.com/kieranrcampbell/ouija as an R package and at http://www.github.com/kieranrcampbell/ouijaflow as a Python/TensorFlow package.

## Introduction

The advent of high-throughput single-cell technologies has revolutionised single-cell biology by allowing dense molecular profiling for studies involving 100-10,000s of cells [1–6]. The increased availability of single-cell data has driven the development of novel analytical methods specifically tailored to single cell properties [7, 8]. The difficulties in conducting genuine time-series experiments at the single-cell level has led to the development of computational techniques known as *pseudotime ordering* algorithms that extract temporal information from snapshot molecular profiles of individual cells. These algorithms exploit studies in which the captured cells behave asynchronously and therefore each is at a different stage of some underlying temporal biological process such as cell differentiation. In sufficient numbers, it is possible to infer an ordering of the cellular profiles that correlates with actual temporal dynamics and these approaches have promoted insights into a number of time-evolving biological systems [9–19].

A predominant feature of current pseudotime algorithms is that they emphasise an “unsupervised” approach. The high-dimensional molecular profiles for each cell are projected on to a reduced dimensional space by using a (non)linear transformation of the molecular features. In this reduced dimensional space, it is hoped that any temporal variation is sufficiently strong to cause the cells to align against a trajectory along which pseudotime can be measured. This approach is therefore subject to a number of analysis choices including gene selection, dimensionality reduction technique, and cell ordering algorithm, all of which could lead to considerable variation in the pseudotime estimates obtained. In order to verify that any specific set of pseudotime estimates are biologically plausible, it is typical for investigators to retrospectively examine specific marker genes or proteins to confirm that the predicted (pseudo)temporal behaviour matches *a priori* beliefs. An iterative “semi-supervised” process maybe therefore be required to concentrate pseudotime algorithms on behaviours that are both consistent with the measured data and compliant with a limited amount of known gene behaviour.

In this paper we present an orthogonal approach implemented within a Bayesian latent variable statistical framework called ‘Ouija’ that learns pseudotimes from small panels of putative or known marker genes (Figure 1A). Our model focuses on switch-like and transient expression behaviour along pseudotime trajectories, explicitly modelling when a gene turns on or off along a trajectory or at which point its expression peaks. Crucially, this allows the pseudotime inference procedure to be understood in terms of descriptive gene regulation events along the trajectory (Figure 1B). As each gene is associated with a particular switch or peak time, it allows us to order the genes along the trajectory as well as the cells and discover which parts of the trajectory are governed by the behaviour of which genes. For example, if the pseudotimes for a set of differentiating cells run from 0 (stem cell like) to 1 (differentiated) and only two genes have switch times less than 0.25 then a researcher would conclude that the beginning of differentiation is regulated by those two genes. We further formulate a Bayesian hypothesis test as to whether a given gene is regulated before another along the pseudotemporal trajectory (Figure 1C) for all pairwise combinations of genes. Furthermore, by using such a probabilistic model we can identify discrete cell types or “metastable states” along continuous developmental trajectories (Figure 1D) that correspond to known cell types.

**Figure 1:**
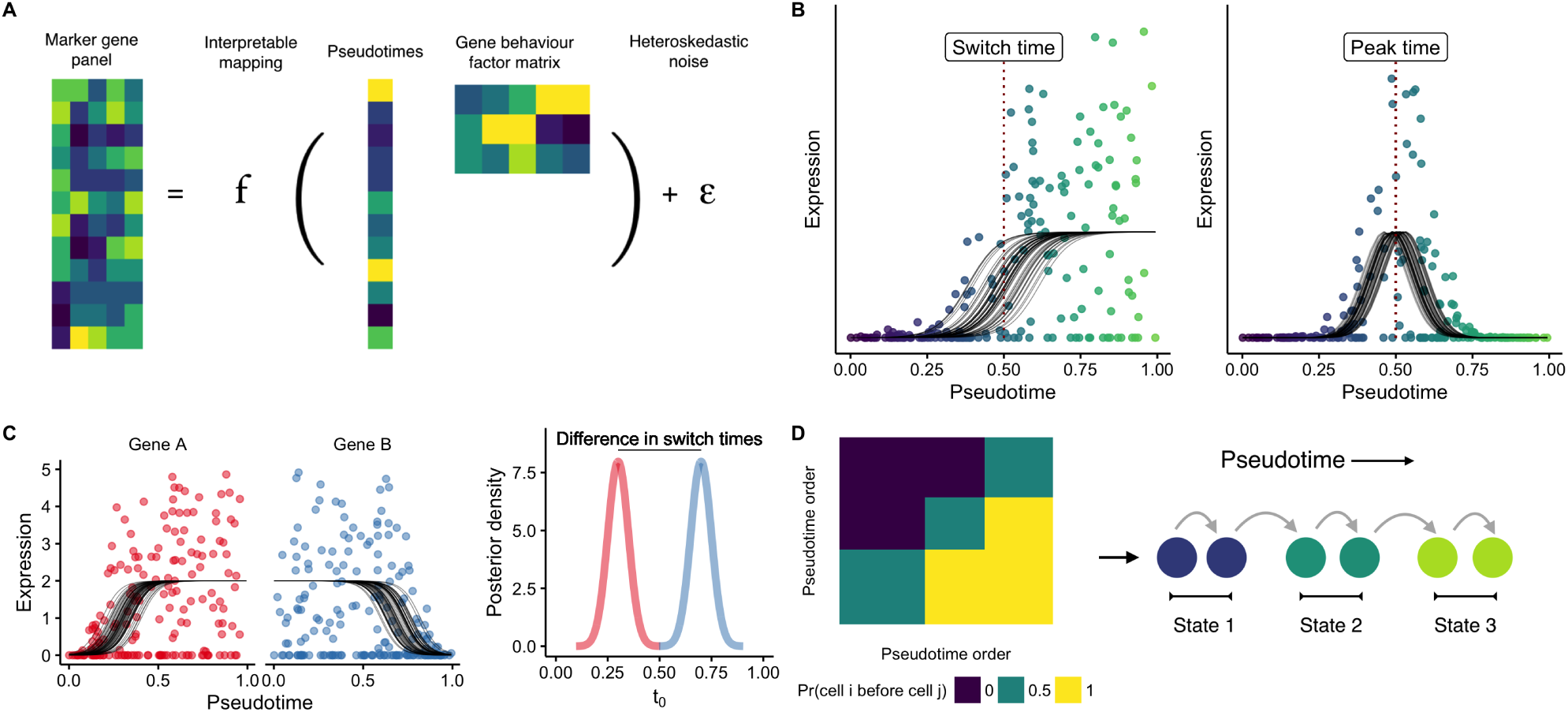
Learning single-cell pseudotimes with parametric models. **A** Ouija infers pseudotimes using Bayesian nonlinear factor analysis by decomposing the input gene expression matrix through a parametric mapping function (sigmoidal or transient). The latent variables become the pseudotimes of the cells while the factor loading matrix is informative of different types of gene behaviour. A heteroskedastic dispersed noise model with dropout is used to accurately model scRNA-seq data. **B** Each gene’s expression over pseudotime is modelled either as a sigmoidal shape (capturing both linear and switch-like behaviour) or through a Gaussian shape (capturing transient expression patterns). These models include several interpretable parameters including the pseudotime at which the gene is switched on and the pseudotime at which a gene peaks. **C** The posterior distributions over the switch and peak times can be inferred leading to a Bayesian statistical test of whether the regulation of a given gene occurs before another in the pseudotemporal trajectory. **D** Ouija can identify discrete cell types that exist along continuous trajectories by clustering the matrix formed by considering the empirical probability one cell is before another in pseudotime.

In the following we introduce our model and demonstrate that it allows pseudotimes equivalent to those inferred using transcriptome-wide models to be learned from only small panel of marker genes. We demonstrate that our model is robust to departures from the prior specification of gene behaviour and that it can identify metastable states along continuous pseudotemporal trajectories consistent with experimentally validated cell types. Finally, we show through simulations that informative Bayesian priors on behaviour parameters may increase the accuracy of pseudotime orderings. An open source implementation of our model is available at http://www.github.com/kieranrcampbell/ouija as an R package and as a Python/TensorFlow package http://www.github.com/kieranrcampbell/ouijaflow.

## Results

### Pseudotime inference from small marker gene panels

The transcriptomes of both single cells and bulk samples exhibit remarkable correlations across genes and transcripts. Such concerted regulation of expression is thought to be due to pathway-dependent transcription [21, 22] and is necessary for the field of network inference from gene expression data [23]. An example of such transcriptome wide correlations can be seen in Figure 2A for the Trapnell et al. [12] dataset, where hierarchical clustering of gene-gene correlations reveals a block-diagonal structure of genes organised into distinct transcriptional pathways. These large correlations across genes imply an intrinsic low-dimensionality of the data meaning it can be efficiently compressed, using techniques such as principal components analysis (PCA). For the Trapnell et al. [12] dataset thousands of genes across the transcriptome exhibit high correlations with the first two principal components (Supplementary Figure 1A) which explain around 20% of the variance (Supplementary Figure 1B).

**Figure 2:**
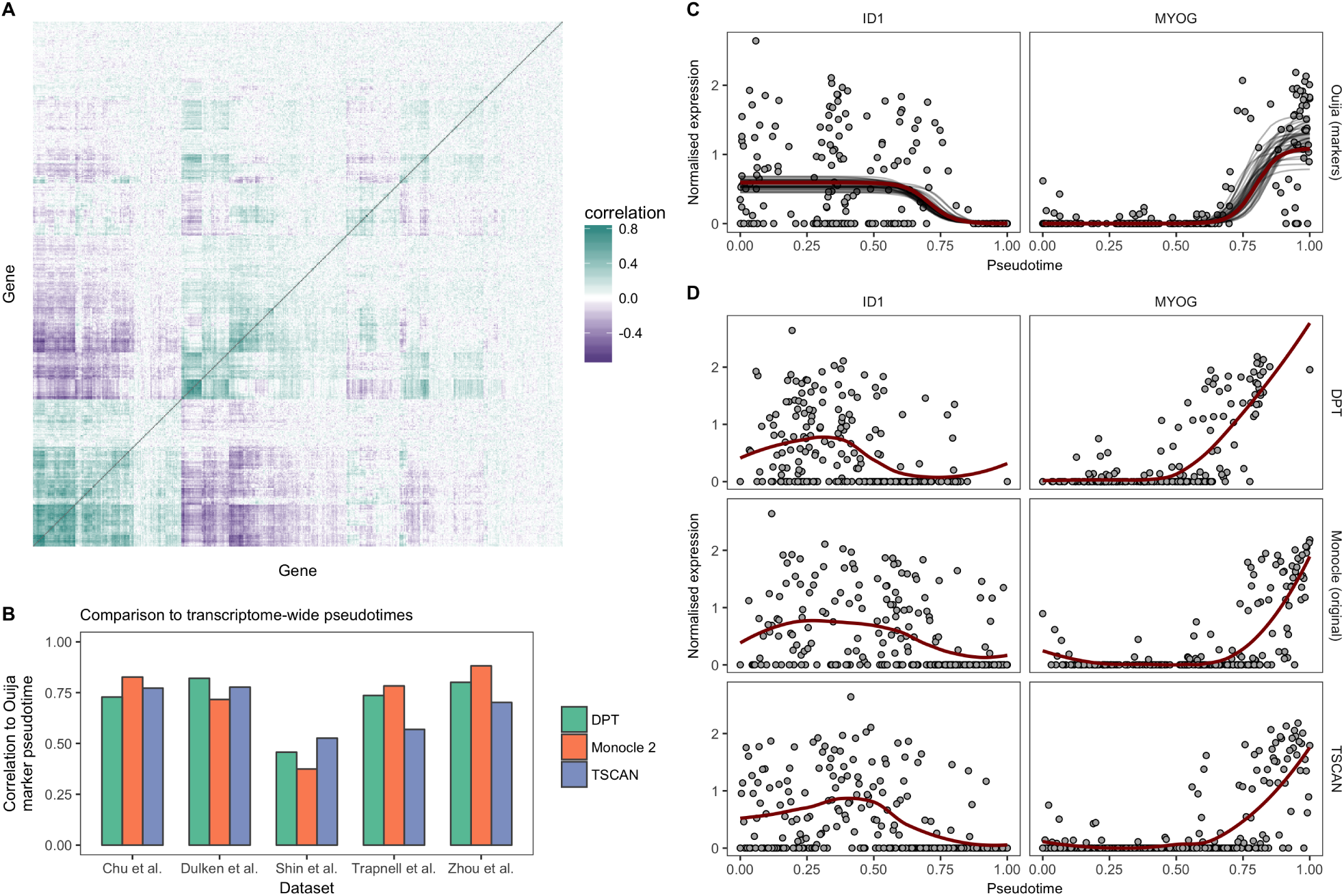
Transcriptome-wide pseudotimes can be inferred from small marker gene panels. **A** A gene-by-gene correlation matrix for the Trapnell et al. [12] dataset reveals similarities in the transcriptional response of hundreds of genes. The redundancy of expression implies the information content of the transcriptome may be compressed through techniques such as principal components analysis (PCA) or by picking informative marker genes. **B** Comparison of pseudotimes fitted using Ouija on a small panel of marker genes to transcriptome-wide fits (using the 500 most variable genes) across five datasets using the algorithms Monocle 2, DPT, and TSCAN. The marker gene fits show high correlation to the transcriptome-wide fits with the exception of the Shin et al. [20] dataset. **C** Gene expression profiles for two marker genes *ID1* and *MYOG* from the Trapnell et al. [12] dataset. The solid red line denotes the maximum *a posteriori* (MAP) Ouija fit while the grey lines show draws from the posterior mean function. **D** Gene expression profiles for the same genes for the algorithms DPT, Monocle 2, and TSCAN show similar expression fits, demonstrating equivalent pseudotemporal trajectories have been inferred. The solid red line denotes a LOESS fit.

This redundancy of expression is often exploited by statistical models of single-cell RNA-seq data. Heimberg et al. [24] use the intrinsic low-dimensionality of the data to reconstruct transcriptome-wide gene expression from ultra-shallow read depths. Cleary et al. [25] apply compressed sensing techniques from the field of signal processing to demonstrate that low-dimensional random projections can efficiently reconstruct high-dimensional gene expression profiles. Rather than explicitly reduce the dimensionality of the data, McCurdy et al. [26] propose a column subset selection procedure whereby a small number of genes are chosen to represent the full transcriptome and demonstrate that this allows clustering of the cells in a similar manner to using the full transcriptomes.

The compressibility of transcriptome data is likewise exploited by many single-cell pseudotime inference algorithms via initial dimensionality reduction steps. For example, Monocle [12] begins by reducing the expression data down to 2 dimensions using independent component analysis, while both TSCAN [18] and Waterfall [16] apply PCA to reduce the data down to 2 dimensions. The implication behind such an approach is that there is sufficient information in just two dimensions of the data via a linear projection to learn “transcriptome-wide” pseudotime and that the majority of expression is redundant given the low-dimensional projection. Nonlinear dimensionality reduction techniques underpin alternative pseudotime approaches such as diffusion components in DPT [27] or reverse graph embedding in Monocle 2 [8].

In Ouija, we exploit the high gene-gene correlations by modelling a small number of *marker* genes that are representative of the whole transcriptome. Such an approach is advantageous as by modelling the data directly rather than a reduced-dimension representation we can understand the pseudotimes for each cell in terms of the behaviour of genes through time rather than abstract notions of manifolds embedded in highdimensional space. This takes the form of a nonlinear factor analysis model, departing from previous models that have relied upon linear factor analysis [28, 29] by introducing sigmoidal nonlinearities (successfully applied previously to single-cell data [30, 31]) and through transient expression functions.

We then turn to the question of how to choose the small number of marker genes in order to fit the pseudotimes. In single-cell pseudotime studies, the cells under examination undergo a known biological process such as differentiation or cell cycle. Importantly, key marker genes associated with these processes are usually known *a priori* by investigators. These marker genes act as positive controls whose behaviour is used post-hoc to confirm the validity of the transcriptome-wide pseudotime fit. For example, in [12] the cells undergo myogenesis and the validity of the pseudotime is confirmed by upregulation of markers of myoblast differentiation such as *MYH3*, *MEF2C*, and *MYOG*, along with down-regulation of markers of actively proliferating cells such as *CDK1* and *ID1*. In [16], the cells undergo neurogenesis and the validity of the transcriptome-wide fit is confirmed via the upregulation of several markers of neural stem cells including *Gfap* and *Sox2*. Further, in [32] the authors tabulate the marker genes they expect to be involved in the process along with their expected behaviour along the differentiation trajectory. Given both the widespread *a priori* knowledge of such markers and their requirement to validate transcriptome-wide pseudotime fits, we therefore propose to derive pseudotimes directly from such markers using our proposed model below.

We first sought to test whether our model applied to small panels of marker genes could accurately recapitulate the transcriptome-wide pseudotimes inferred by popular pseudotime methods. We applied Monocle 2, DPT, and TSCAN to five publicly available single-cell RNA-seq datasets [12, 16, 33–35] using the 500 most variable genes as input (the default in packages such as Scater [36] for PCA representations). For each dataset, we then inferred pseudotimes using Ouija based only on a small number of marker genes reported in each paper (ranging from 5 to 12), and compared the Pearson correlation between the Ouija pseudotimes and the pseudotimes reported for each dataset (Figure 2B). There was good agreement between the marker-based pseudotimes inferred using Ouija and the transcriptome-wide pseudotimes inferred using existing algorithms, with the correlation exceeding 0.75 in the majority of comparisons.

Noting that the correlation will not be 1 unless the algorithms are identical, we sought to compare Ouija’s correlation to transcriptome-wide pseudotime to the agreement of the transcriptome-wide pseudotimes with each other. We found large variability in the agreement between existing algorithms using transcriptome-wide pseudotimes, with correlations as high as 0.93 but as low as 0.61 (Supplementary Figure 2). We found the marker-based Ouija pseudotimes have higher correlations to one of the transcriptome-wide algorithms than they have amongst each other in all but one of the datasets studied. On average, the correlation between Ouija’s marker based pseudotime with the transcriptome-wide pseudotimes was around 0.1 lower than the correlation amongst the transcriptome-wide pseudotimes, though given Ouija uses around 1-2% the number of input genes we believe this is a positive result that represents transcriptome-wide pseudotimes may be inferred using interpretable, parametric models on a small number of marker genes chosen *a priori*.

This equivalence of transcriptome-wide and marker-based pseudotimes is further confirmed by examining the qualitative fit of the marker genes across the different algorithms. For example, Figure 2C shows the posterior fit of the marker-based pseudotime for two marker genes from [12], correctly inferring the switch-like downregulation of *ID1* and the upregulation of *MYOG*. Near identical behaviour is found using transcriptome-wide pseudotimes derived from DPT, Monocle, and TSCAN (Figure 2D). We note the low correlations of the marker-based Ouija pseudotimes with the transcriptome-wide fits for the Shin et al. dataset. Upon close inspection of the marker genes (Supplementary Figure 3) we found that the expression of four of the marker genes (*Aldoc*, *Apoe*, *Eomes*, *Sox11*) were highly correlated (the switch times are similar) whilst *Gfap* and *Stmn1* showed little variation over pseudotime. This meant that there was effectively only a single marker gene for this data set - too few for reliable marker gene-based pseudotime inference.

### Gene regulation timing from marker gene-based pseudotime

Having demonstrated Ouija can accurately recapitulate transcriptome-wide pseudotimes using just small marker gene panels, we next sought to show how it allows for marker-driven inference of such trajectories. Most pseudotime inference algorithms (such as Monocle 2, DPT, TSCAN, Slicer [37]) emphasise that cells occupy a low-dimensional manifold embedded in high-dimensional space and traversing this manifold corresponds to following the cells over pseudotime. While such an approach is theoretically well grounded it is difficult to understand *why* the procedures result in a particular pseudotime trajectory, leading to the post-hoc marker gene examination procedures discussed above as validation and to add interpretability.

To demonstrate that Ouija allows for feature-driven inference of single-cell pseudotime we applied it to a single-cell time-series dataset of human embryonic stem cells differentiating into definitive endoderm cells. The authors examined the expression of key marker genes over time and found 9 to exhibit approximately switch-like behaviour (*POU5F1*, *NANOG*, *SOX2*, *EOMES*, *CER1*, *GATA4*, *DKK4*, *MYCT1*, and *PRDM1*) with a further two exhibiting transient expression (*CDX1* and *MSX2*). We applied Ouija using noninformative priors over the behaviour parameters with no information about the capture times of the cells included.

The resulting pseudotime fit demonstrates we can understand single-cell pseudotime in terms of the behaviour of particular genes. Figure 3A shows a heatmap of the 9 switch-like genes (top) and 2 transient genes (bottom), ordered by the posterior switch time of each gene. It can be seen that the early trajectory is characterised by the expression of *NANOG*, *SOX2*, and *POUF51*, which then leads to a cascade of switch-like activation of the remaining genes as the cells differentiate.

**Figure 3:**
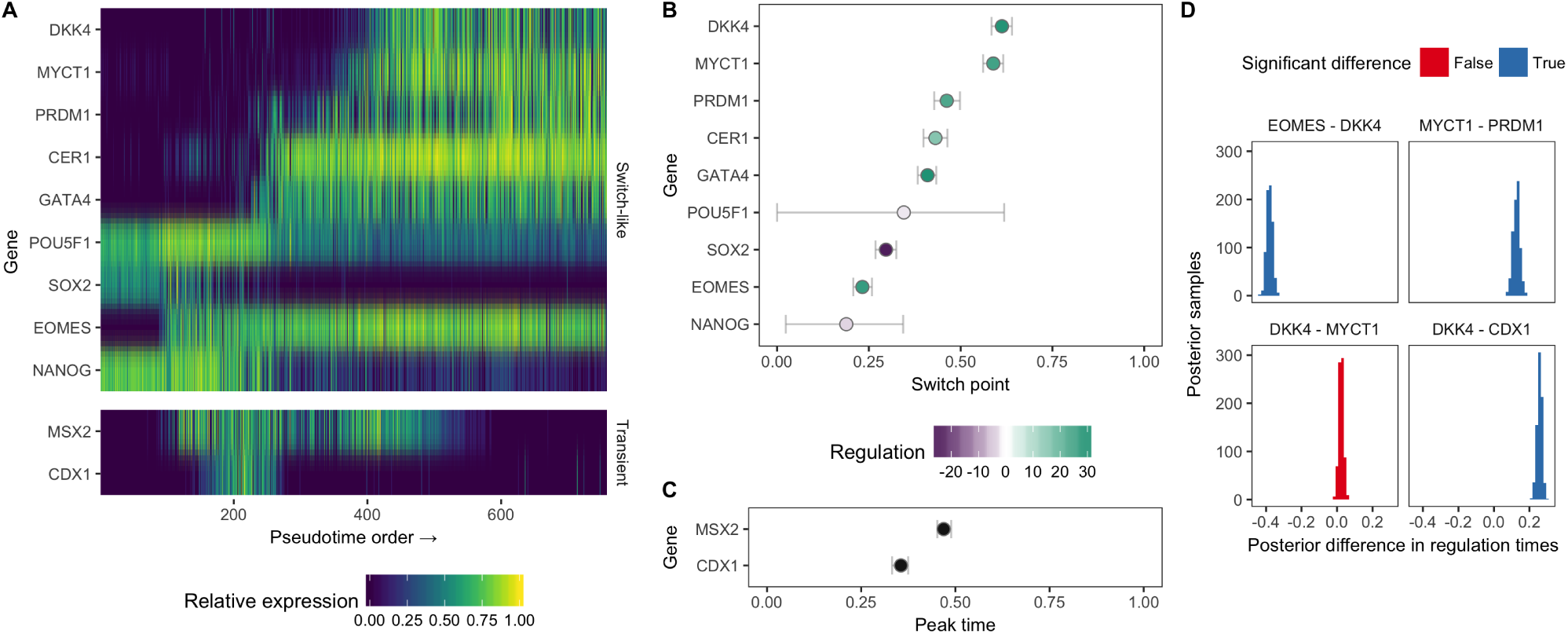
Parametric models lead to pseudotimes centred around gene regulation timing. **A** An expression heatmap for the 9 switch-like genes and 2 transient genes in the Chu et al. dataset, with genes ordered by the posterior mean of the switch time. **B-C** Posterior distributions over the switch times and peak times for the 11 genes, coloured by their up or down regulation along pseudotime. The horizontal error bars show the 95% highest probability density credible intervals. **D** A Bayesian hypothesis test can quantify whether the posterior difference between two regulation timings (either switch or peak time) is significantly different from 0, allowing us to determine whether a given gene is regulated before or after another along pseudotime.

While transcriptome-wide pseudotime algorithms could provide similar heatmaps if the marker genes were known in advance, the key departure of Ouija is that we can quantitatively associate each gene with a region of pseudotime at which its regulation (switch time or peak time) occurs. This is illustrated in Figure 3B-C showing the posterior values for the regulation timing along with the associated uncertainty. In essence, Ouija allows us to order *genes* along trajectories as well as being able to order the cells, which provides insight into gene regulation relationships.

To approach such questions of gene regulation timings in a quantitative and rigorous manner we constructed a Bayesian hypothesis test to find out whether one gene is regulated before another given the noise in the data. If 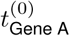 and 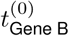 are the regulation timings of genes A and B respectively, we calculate the posterior distribution 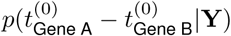, and if both the lower and upper bounds of the 95% posterior credible interval fall outside 0 we say the two genes are regulated at significantly different times. We applied this to the pseudotime fit in the Chu *et. al.* dataset, the results of which can be seen in Figure 3D for a subset of genes. The model suggests that *EOMES* is downregulated before *DKK4* and *MYCT1* is downregulated after *PRDM1*. Furthermore, it suggests the switch-like downregulation of *DKK4* occurs after the transient peak-time of *CDX1*. However, it suggests the difference in regulation timings of *DKK4* and *MYCT1* are not significantly different from zero, which could imply co-regulation.

### Ouija is robust to gene behaviour misspecification

A potential disadvantage of our model is the requirement to pre-specify genes as having switch-like or transient behaviour over pseudotime, which may result in biased or erroneous pseudotimes. We noticed such an effect in the Li *et al.* (2016) [32] dataset, where the authors pre-specified how they expected several marker genes to behave over pseudotime. Upon fitting the pseudotimes using Ouija, we noted that the genes *Mef2c* and *Pik3r2* exhibited the correct upregulation over pseudotime (Figure 4A), but that *Scd1* that was supposed to exhibit transient, peaking expression was effectively constant along the trajectory (Figure 4B).

**Figure 4:**
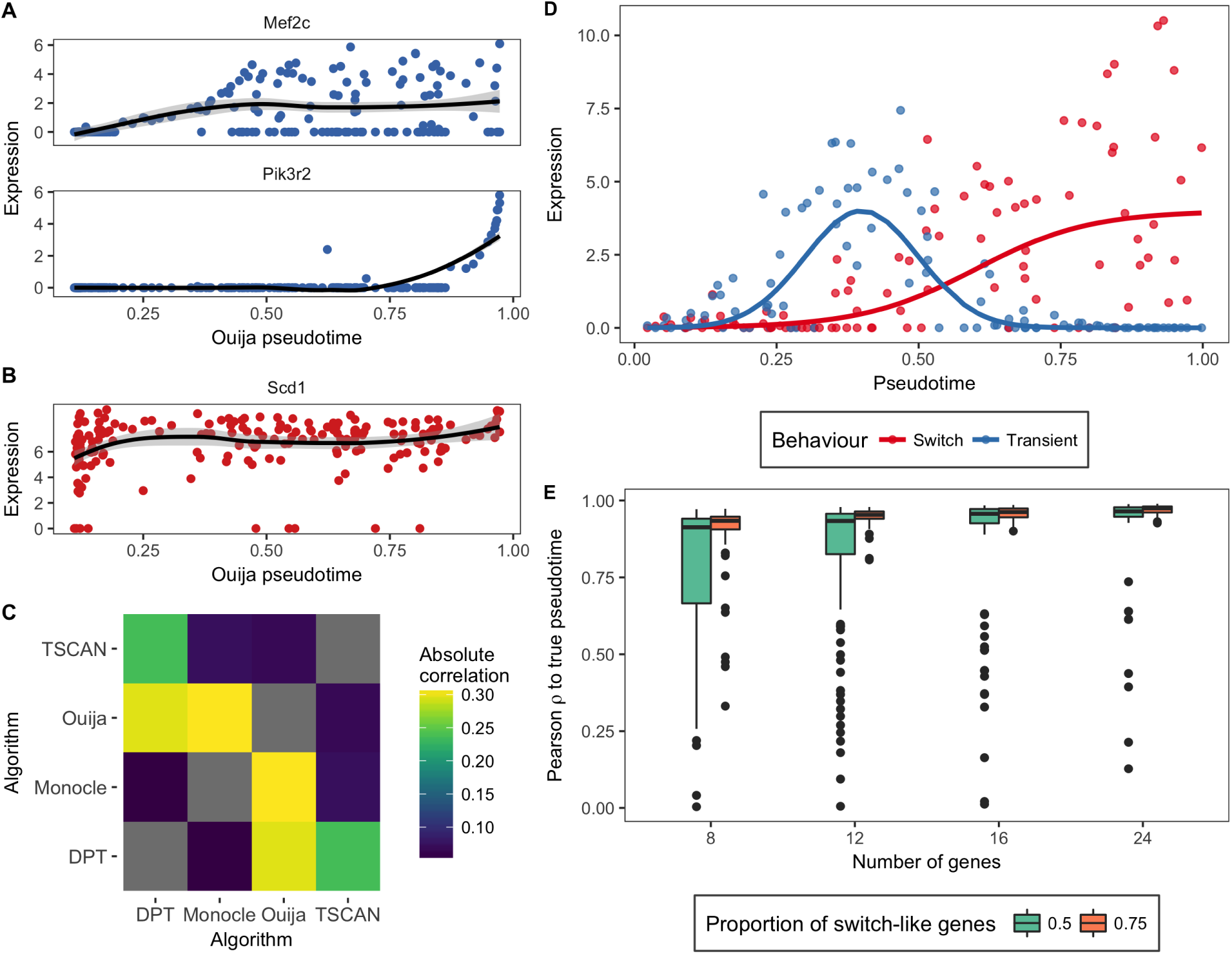
Ouija is robust to gene behaviour misspecification. **A** The genes *Mef2c* and *Pik3r2* show the expected behaviour in a marker-based pseudotime fitted to the Li *et al.* (2016) [32] dataset (“constant upregulation” and “transient upregulation” respectively). However, the gene *Scd1* (**B**) was claimed to have “tide wave” regulation (transient expression), but a LOESS fit over pseudotime (black line) shows effectively constant expression over pseudotime. **C** We found very low agreement between the different pseudotime inference algorithms for this dataset. Curiously, the largest agreement was reported between Ouija using only markers and Monocle 2 using the 500 most variable genes. **D** We simulated datasets with genes either exhibiting switch-like expression over pseudotime or transient expression, with an overdispersed, zero-inflated noise model to mimic real data. **E** Ouija was benchmarked assuming all genes were switch-like when a certain proportion were actually transient across a range of geneset sizes. Even at only 8 genes, half of which are actually transient, Ouija still recovers a median correlation of greater than 0.9 with the true pseudotime, which only increases with increasing number of genes and switch-like behaviour.

We first sought to discover whether this was a particular failing of Ouija or a result common to all pseudotime algorithms. To do so, we fitted transcriptome-wide pseudotimes using TSCAN, Monocle 2 and DPT, and compared both the correlation among the different algorithms and the behaviour of the specified marker genes. We found remarkably low correlations between the different pseudotime algorithms (Figure 4C), with the highest correlations reported between Ouija using markers only and Monocle 2 using the full transcriptome. Furthermore, none of the pseudotime fits displays consistent nor expected behaviour for the set of marker genes (Supplementary Figure 4). For example, the gene *Foxa2* is seemingly dowregulated under DPT, upregulated under Monocle 2 and Ouija, and exhibits transient expression under TSCAN.

Next, we performed extensive simulations to discover the extent to which Ouija is in general robust to gene behaviour misspecification. We simulated datasets where either 75% or 50% of the genes were switch-like (Figure 4D) for 8, 12, 16 & 24 genes with 100 replications for each situation, and re-inferred the pseudotimes using Ouija assuming all genes were switch-like. The results can be seen in Figure 4D. Even with 4 switch-like and 4 transient genes Ouija still achieves a median correlation greater than 0.9 with the true pseudotimes, a result that only increases with more switch-like genes. We believe this shows that Ouija is highly robust to misspecification of prior knowledge of gene behaviour.

It is further possible to identify errors in the prior belief of gene behaviour without having to explicitly fit a pseudotemporal trajectory. If a dataset contains a number of switch-like and transient genes, the switch-like genes will have high absolute correlation with themselves but low absolute correlation with the transient genes, which will in turn have high absolute correlation with themselves. This effect is exemplified in the Chu *et al.* dataset that contains 9 switch-like and 2 transient genes. A hierarchical clustering of the absolute correlations across the genes reveals the transient genes clustering separately from the switch-like genes (Supplementary Figure 5). Therefore, an investigator could corroborate their prior expectations through similar investigations.

### Identifying discrete cell types along continuous developmental trajectories

We further investigated the single cell expression data from a study tracking the differentiation of embryonic precursor cells into haematopoietic stem cells (HSCs) [33]. The cells begin as haemogenic endothelial cells (ECs) before successively transforming into pre-HSC and finally HSC cells. The authors identified six marker genes that would be down-regulated along the differentiation trajectory, with early down-regulation of *Nrp2* and *Nr2f2* as the cells transform from ECs into pre-HSCs, and late down-regulation of *Nrp1*, *Hey1*, *Efnb2* and *Ephb4* as the cells emerge from pre-HSCs to become HSCs. The study investigated a number of distinct cell types at different stages of differentiation: EC cells, T1 cells (*CDK*45^*−*^ pre-HSCs), T2 cells (*CDK*45^+^ pre-HSCs) and HSC cells at the E12 and E14 developmental stages.

We therefore sought to identify the existence of these discrete cell types along the continuous developmental trajectory. As Ouija uses a probabilistic model and inference we were able to obtain a posterior ordering “consistency” matrix (Figure 5A) where an entry in row *i* column *j* denotes the empirical probability that cell *i* is ordered before cell *j*. Performing PCA on this matrix gives a rank-one representation of cell-cell continuity, which is then clustered using a Gaussian mixture model to find discrete cell states along the continuous trajectory (where the number of states is chosen such that the Bayesian information criterion is maximised).

**Figure 5:**
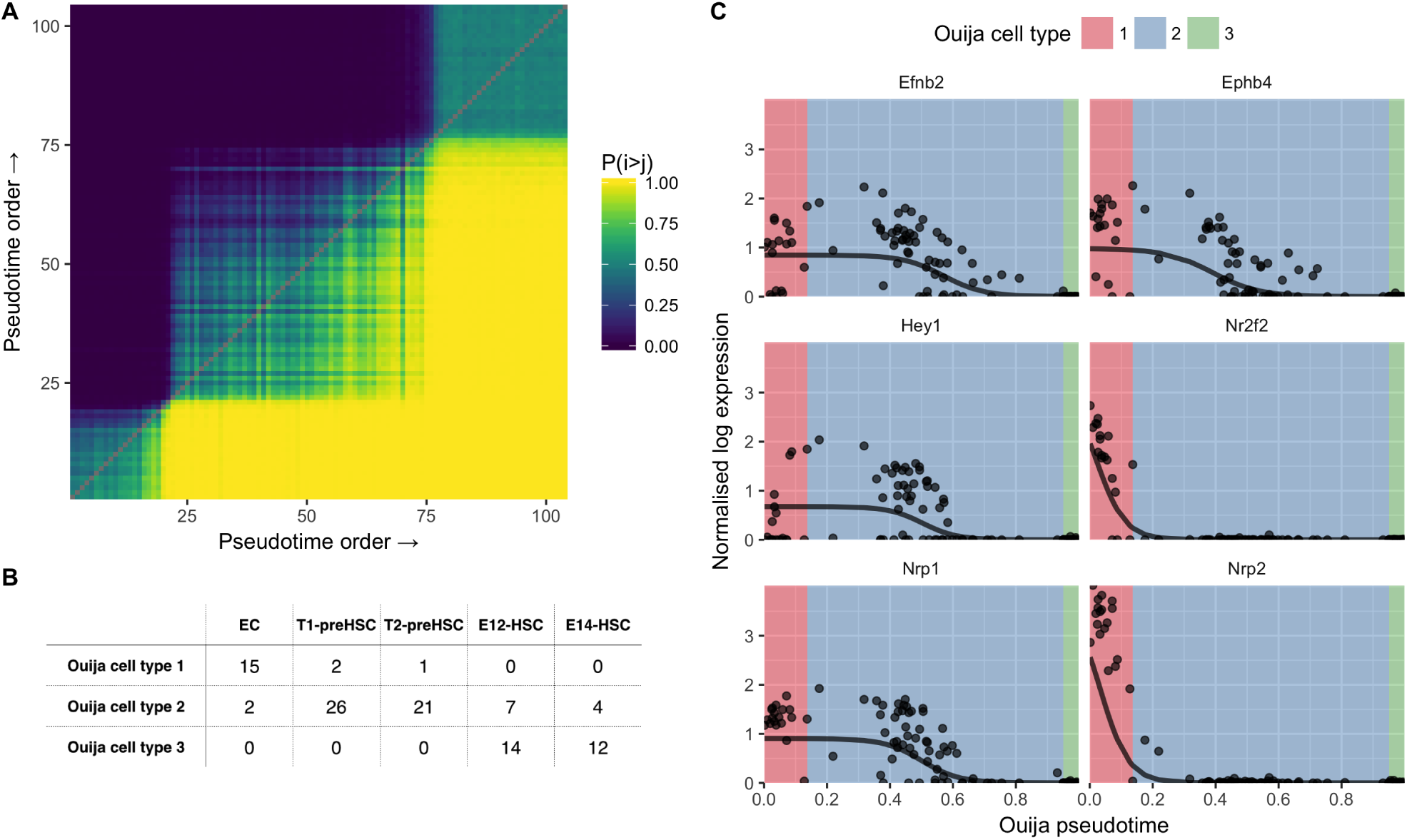
Pseudotime ordering and cell type identification of haematopoeietic stem cell differentiation. **A** Consistency matrix of pseudotime ordering. Entry in the *i*^*th*^ row and *j*^*th*^ column is the proportion of times cell *i* was ordered before cell *j* in the MCMC posterior traces. Gaussian mixture modelling on the first principal component of the matrix identified three clusters that are evident in the heatmap. **B** Confusion matrix for cell types identified in original study (columns) and Ouija inferred (rows). Ouija inferred cluster 1 largely corresponds to EC cells, cluster 2 corresponds to pre-HSC cells while cluster 3 corresponds to HSC cells. **C** HSC gene expression as a function of pseudotime ordering for six marker genes. Background colour denotes the maximum likelihood estimate for the Ouija inferred cell type in that region of pseudotime.

Applying this methodology to the Zhou *et. al.* dataset uncovered three metastable groups of cells corresponding to endothelial, pre-HSCs and HSCs respectively (Figure. 5B). Misclassifications within cell types (T1/T2 and E12/E14 cells) could be explained by examining a principal components analysis of the global expression profiles (Supplementary Figure 6) which suggests that these cell types are not completely distinct in terms of expression. When examining the inferred pseudotime progression of each marker gene (Figure 5C), these three metastable states corresponded to the activation of all genes at the beginning of pseudotime time, the complete inactivation of all the marker genes at the end of the pseudotime and a intervening transitory period as each marker gene turns off. Each metastable state clearly associates with a particular cell type with *Nrp2* and *Nr2f2* exhibiting early down-regulation and *Nrp1*, *Hey1*, *Efnb2* and *Ephb4* all exhibiting late down-regulation. Using this HSC formation system as a proof-of-principle it is evident that, if a small number of switch-like marker genes are known, it is possible to recover signatures of temporal progression using Ouija and that these trajectories are compatible with real biology.

To show the widespread applicability of this method we applied it to two further publically available datasets. Dulken *et. al.* [34] examined the trajectory of quiescent neural stem cells (qNSCs) as they differentiate into activated neural stem cells (aNSCs) and neural progenitor cells (NPCs). Applying Ouija’s clustering-along-pseudotime revealed seven distinct clusters (Supplementary Figure 7; Supplementary Table 1) with clusters 1-2 corresponding to early and late qNSCs, cluster 3 defining the qNSC to aNSC transition, clusters 4-6 corresponding to early to late aNSCs and cluster 7 defining the aNSC to NPC transition. We similarly applied this method to the Chu et al. dataset of time-series scRNA-seq that identified 8 distinct clusters along pseudotime (Supplementary Figure 8; Supplementary Table 2). Clusters 1-4 track the cells as the progress through the 4 stages from 0 hours to 36 hours, while clusters 5-8 track the 3 stages from 36 hours to 96h hours but with much more heterogeneity within each cluster, which is expected due to the longer time-scales considered.

### Incorporating prior information can improve pseudotime inference

A particular advantage of using Bayesian models with interpretable parameters is that we may express any prior knowledge about the gene behaviour as informative priors. For example, for each gene we model as switch-like there is the switch strength parameter *k* that models how quickly a gene is upregulated if *k* is positive or how quickly it is downregulated if it is negative. A researcher may have a firm prior belief that a gene will be up or downregulated along the trajectory and thus can place a prior *p*(*k*) on the particular parameters. Using Bayes’ rule, the posterior distribution of both the pseudotimes and gene-specific parameters is then calculated by combining this informative prior with the data likelihood. The crucial observation here is that the posterior distribution of the pseudotimes is affected by priors on the gene behaviour parameters, meaning incorporating prior information about gene behaviours may improve pseudotime inference. Such informative priors may be placed on any of the parameters that govern interpretable gene behaviour. For example, if a researcher expects a particular transient gene to peak early in the trajectory then they may encode this using a prior distribution on the peak time.

We sought to test the extent to which incorporating knowledge of gene behaviours through informative Bayesian priors aids pseudotime inference. To do so we performed extensive simulations of single-cell pseudotime under monotonic changes in expression and reinferred using Ouija with both noninformative and informative priors, as well as DPT and TSCAN. In order to emulate the fact that the data will not truly come from a sigmoidal link function, we simulated data from various link functions used in logistic regression including probit and complementary log-log (Figure 6) along with a “threshold” model where the expression is on or off with a particular probability that changes along the trajectory (see supplementary text for full details).

**Figure 6:**
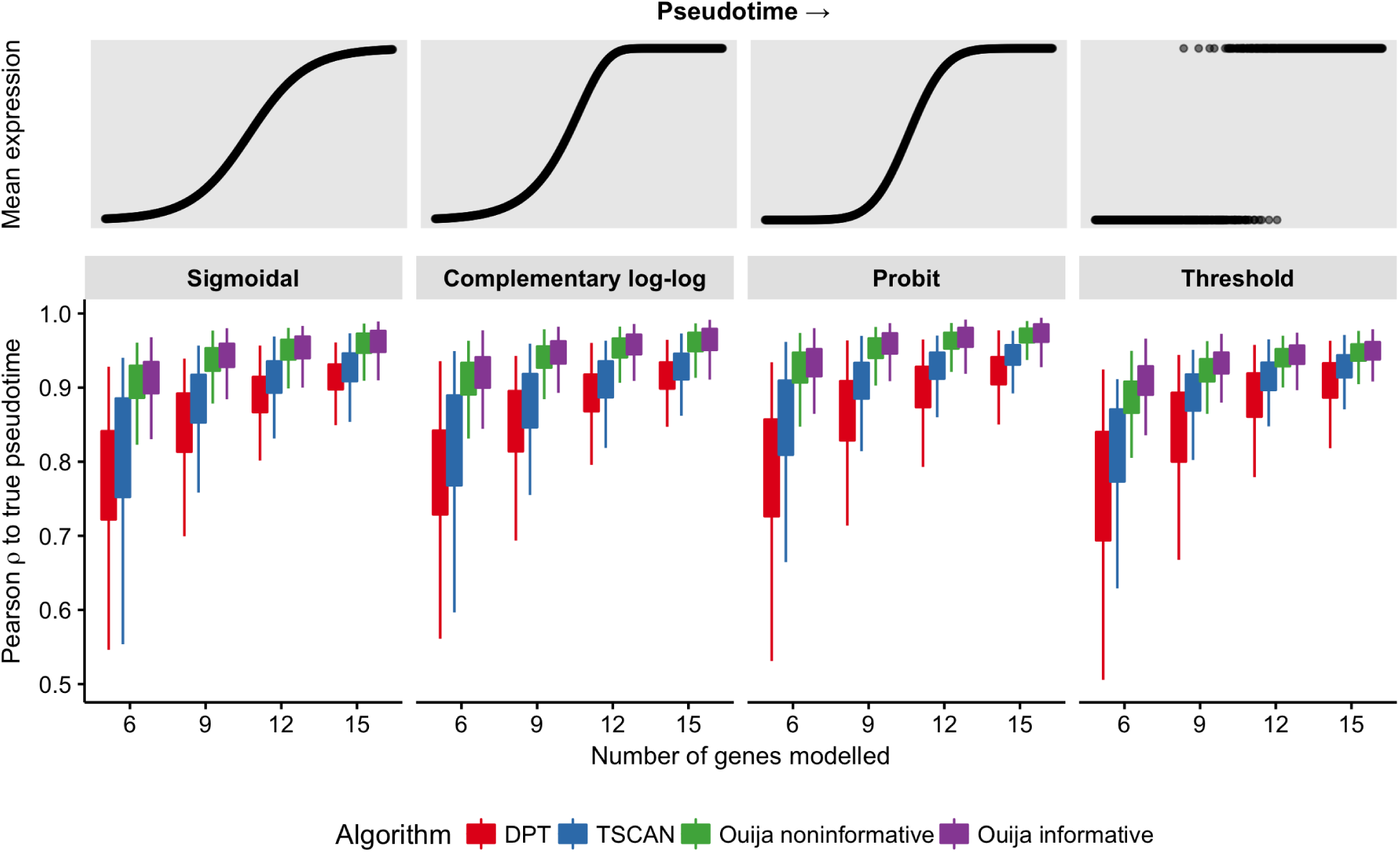
Incorporating prior information can improve pseudotime inference. We sought to identify the benefits of incorporating prior information about the behaviour of genes to the accuracy of pseudotime inference. We simulated data according to four different mean functions (sigmoidal, complementary log-log, probit, and threshold) under identical noise model and reinferred using DPT, TSCAN, Ouija with noninformative priors, and Ouija with informative priors. The results show a marginal though significant gain in inference when incorporating prior knowledge.

The results can be seen in Figure 6, with similar characteristics across the four mean functions considered. In all cases Ouija performs substatially better than DPT and TSCAN, but we note that this is likely due to the data generating model more closely matching the likelihood model of Ouija though could also be explained by the fact that DPT and TSCAN are not designed for small panels of genes. In each case the gain from incorporating prior information is statistically significant (Supplementary Table 3), but we note that the effect sizes are in practice quite small. Since to infer a consistent pseudotime, sufficient correlations must exist in the data, prior knowledge may only make a relatively minor contribution. However, researchers dealing with data with low biological signal to noise ratio may find it advantageous to incorporate such constraints to improve the quality of their inferences.

### Scalable pseudotime inference using TensorFlow

Finally, we wanted to consider a study composed of a large panel of putative marker genes to determine if Ouija could automatically identify genes satisfying its behavioural constraints. We identified a single-cell RNA-seq study [38] that examined variation between individual hematopoietic stem and progenitor cells from two mouse strains (C57BL/6 and DBA/2) as they age. Principal component analysis for each cell type and age showed a striking association of the top principal components with cell cycle-related genes (Figure 7A), indicating that transcriptional heterogeneity was dominated by cell cycle status. They scored each cell for its likely cell cycle phase using signatures based on functional annotations [39] and profiles from synchronized HeLa cells [40] for the G1/S, S, G2, and G2/M phases.

**Figure 7:**
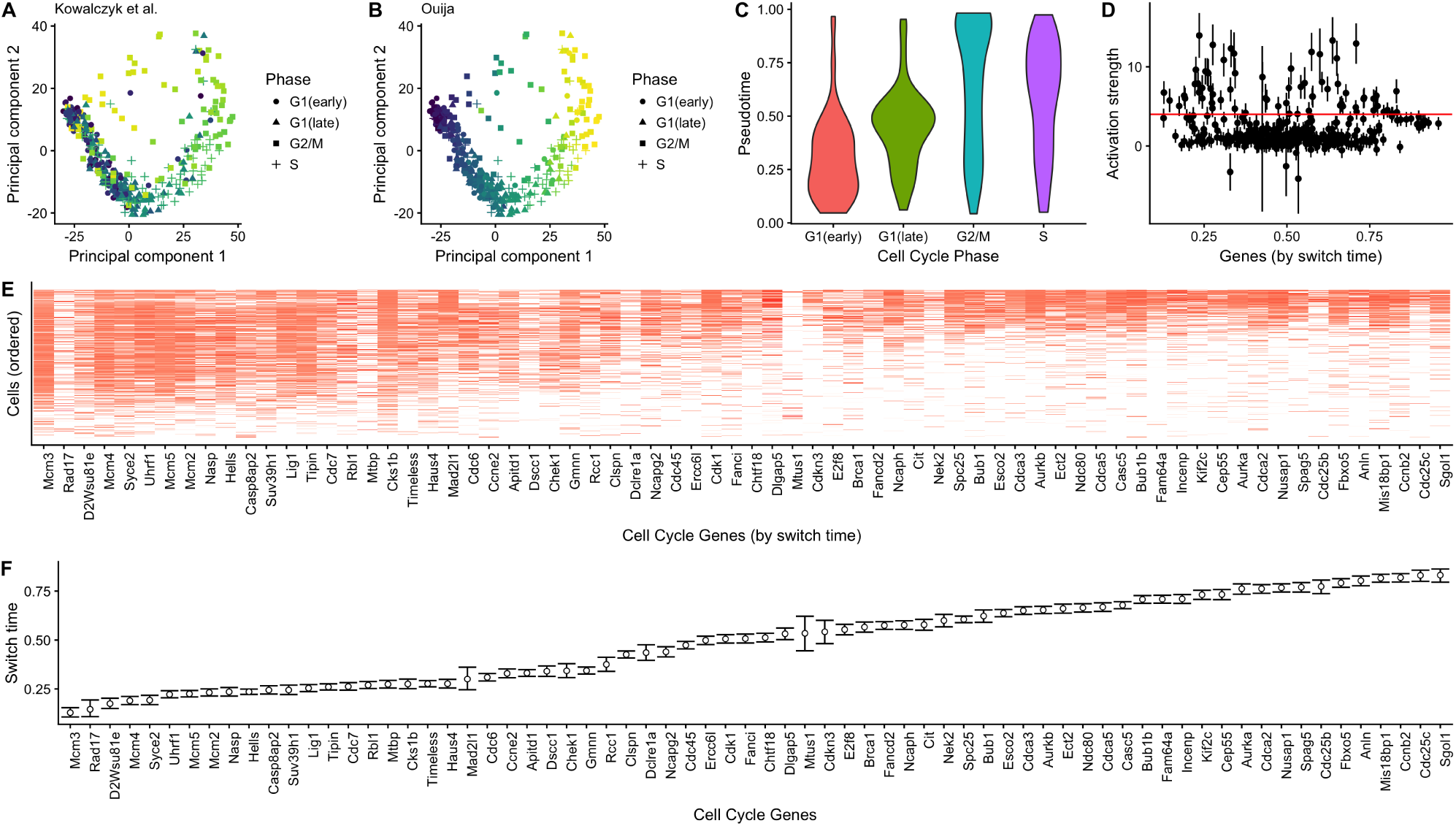
Cell cycle phase prediction. Principal component representation of hematopoietic stem cells coloured according to (A) the original cell cycle progression score [38] and (B) Ouija - cell cycle classes indicated are based on original study classifications. (C) Distribution of Ouija inferred pseudotime versus the original cell cycle classifications. (D) Estimated activation strengths for the 374 cell cycle gene panels. (E) Gene expression profile for 88 switch-like genes with cells ordered by pseudotime and (F) genes ordered by activation time.

We investigated if Ouija could be used to identify cell cycle phase, treating the inferential problem as a continuous pseudotime process and assuming all genes as candidate switch genes. We applied Ouija to 1,008 C57Bl/6 HSCs using 374 GO cell cycle genes that satisfied gene selection criteria used in the original study. This large number of genes and cells makes inference using Hamiltonian Monte Carlo (HMC) slow so we implemented a second version of Ouija (termed *Ouijaflow*) using the probabilistic programming language Edward [41] based on TensorFlow [42]. This performs fast approximate Bayesian inference using reparametrization gradient variational inference.

The estimated pseudotime progression given by Ouija recapitulates the trajectory observed in principal component space (Figure 7A). The estimated pseudotime distribution correlates well with the cell cycle phase categorisation given in the original study (Figure 7C). Furthermore, we identified 88 genes with large activation strengths indicating strong switching-on behaviour (Figure 7D). Ordering the genes by activation time demonstrates a cascade of expression activation across these 88 genes over cell cycle progression with the quiescent (*G*_0_) indicated by complete inactivation of all 88 genes (Figure 7E,F). The explicit parametric model assumed by Ouija makes this gene selection and ordering process simple and *quantitative* compared to a non-parametric approach that would require some retrospective analysis or visual inspection.

## Conclusions

We have developed a novel approach for pseudotime estimation based on modelling switch-like and transient expression behaviour for a small panel of marker genes chosen *a priori*. Our strategy provides an orthogonal and complimentary approach to unsupervised whole-transcriptome methods that do not explicitly model any gene-specific behaviours and do not readily permit the inclusion of prior knowledge.

We demonstrate that the selection of a few marker genes allows comparable pseudotime estimates to whole transcriptome methods on real single cell data sets. Furthermore, using a parametric gene behaviour model and full Bayesian inference we are able to recover posterior uncertainty information about key parameters, such as the gene activation time, allowing us to explicitly determine a potential ordering of gene (de)activation and peaking events over (pseudo)time. The posterior ordering uncertainty can also be used to identify homogeneous metastable phases of transcriptional activity that might correspond to transient, but discrete, cell states.

Furthermore, whilst we do not explicitly address branching processes in this work, our framework provides a natural and simple extension to allow for multiple lineages and cell fates using a sparse mixture of factor analyzers in which each lineage is denoted by a separate mixture component and the factors loadings are shrunk to common values to denote shared branches. This has been explored in work elsewhere [43].

In summary, Ouija provides a novel contribution to the increasing plethora of pseudotime estimation methods available for single cell gene expression data.

## Methods

### Overview

We give a high-level overview of our pseudotime inference framework here and provide more technical details in the following sub-sections. The aim of pseudotime ordering is to associate a *p*-dimensional expression measurement (the data) to a latent unobserved pseudotime. Mathematically we can express this as the following:

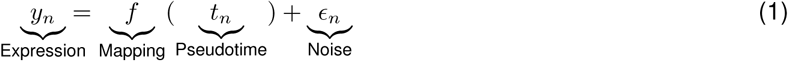

where the function *f* maps the one-dimensional pseudotime *t*_*n*_ for cell *n* to the *p*-dimensional observation space in which the data lies. The challenge lies in the fact that *both* the mapping function *f* and the pseudotimes are *unknown*. Our objective here is to use parametric forms for the mapping function *f* that will enable relatively fast computations whilst characterising certain gene expression temporal behaviours. The specification of a statistical pseudotime algorithm therefore comes down to the choice of the mean function *f* and the noise model on *ε* that we detail below.

The approach we adopt is therefore a form of latent variable model implemented as *non-linear parametric factor analysis* where the factors correspond to the pseudo-times and the factor loadings correspond to the interpretable parameters of the sigmoidal or transient mean functions that provide the non-linearity. In addition, we model dropouts and a strict empirically motivated mean-variance relationship which is required to provide constraints on the latent variable model since nothing on the right hand side of equation 1 is actually measured or observed.

### Statistical model

Input data normalisation

We index *N* cells by *n* ∈ 1*,…, N* and *G* genes by *g* ∈ 1*,…, G*. Let *y*_*ng*_ = [Y]_*ng*_ denote the log-transformed non-negative observed cell-by-gene expression matrix. In order to make the strength parameters comparable between genes we normalise the gene expression so the approximate half-peak expression is 1 through the transformation

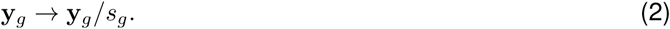

where *s*_*g*_ is a gene-specific size factor defined by

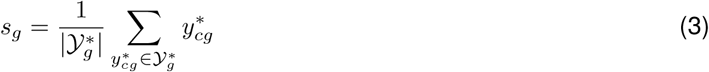

and 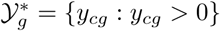.

### Noise model

Our statistical model can be specified as a Bayesian hierarchical model where the likelihood is given by a bimodal distribution formed from a mixture of zero-component (dropout) and an non-zero expressing cell population. If *µ*(*t_n_,* Θ_*g*_) is the mean for cell *n* and gene *g* (evaluated at pseudotime *t*_*n*_ with gene-specific parameters Θ_*g*_) then

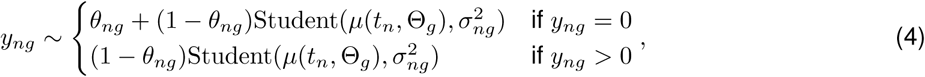

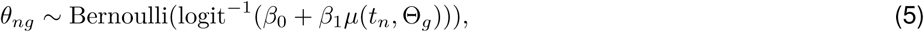

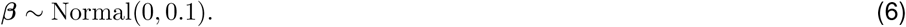

The relationship between dropout rate and expression level is expressed as a logistic regression model [44]. Furthermore, we impose a mean-variance relationship of the form:

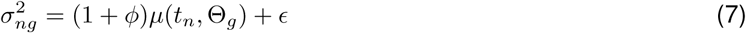

where *ϕ* is the dispersion parameter with prior *ϕ ~* Gamma(*α_ϕ_, β*_*ϕ*_), which is motivated by empirical observations of marker gene behaviour (see supplementary text).

### Mean functions

We then need to specify the form of the mean functions *µ*(*t_n_,* Θ_*g*_), for which we consider both sigmoidal and transient genes.

For the sigmoidal case we have

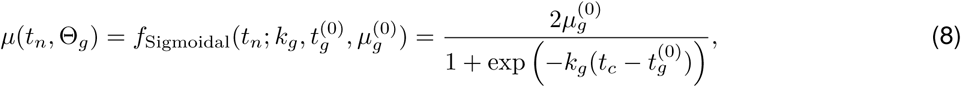

where *k*_*g*_ and 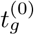 denote the activation strength and activation time parameters for each gene and 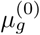 the average peak expression with default priors

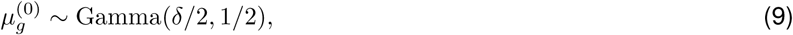

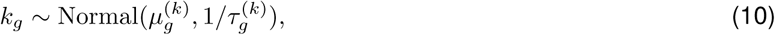

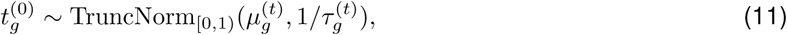

If available, user-supplied prior beliefs can be encoded in these priors by specifying the parameters 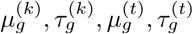.Otherwise, inference can be performed using uninformative hyperpriors on these parameters. Specifying 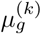 encodes a prior belief in the strength and direction of the activation of gene *g* along the trajectory with 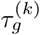 (inversely-) representing the strength of this belief. Similarly, specifying 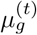 encodes a prior belief of where in the trajectory gene *g* exhibits behaviour (either turning on or off) with 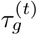 encoding the strength of this belief.

For the transient case we have

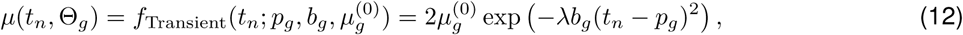

where we take *λ* to be a constant *λ* = 10 and with a prior structure

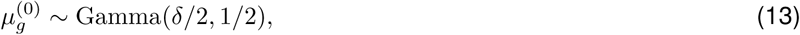

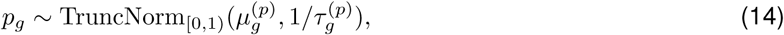

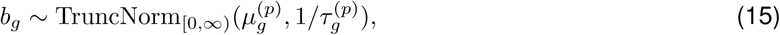

where informative priors may be placed on *p* and *b* as before.

Note that if the mapping functions *f* are restricted to a linear form then the model reduces to Factor Analysis. In other words, performing factor analysis on single-cell RNA-seq data is entirely equivalent to finding a trajectory where gene expression is linear over time with no prior expectations on how the genes behave. If we model a common precision across all genes so *τ*_*g*_ = *τ* then then model reduces further to probabilistic principal components analysis [45], providing an explicit interpretation for the results of principal component analysis on single-cell data.

### Inference

We performed posterior inference using Markov Chain Monte Carlo (MCMC) stochastic simulation algorithms, specifically the No U-Turn Hamiltonian Monte Carlo approach [46] implemented in the STAN probabilistic programming language [47] which we use to implement our model. The parameter *ε* = 0.01 is used to avoid numerical issues in MCMC computation. For larger marker gene panels, such as in the cell cycle analysis section, we used the reparametrization gradient variational inference methods implemented in Edward [41] to perform approximate Bayesian inference.

## Author’s contributions

K.R.C. and C.Y. conceived the study. K.R.C. developed software and performed computer simulations. K.R.C. and C.Y. wrote the manuscript.

## Acknowledgements

K.R.C. is supported by a UK Medical Research Council funded doctoral studentship. C.Y. is supported by a UK Medical Research Council New Investigator Research Grant (Ref. No. MR/L001411/1), the Wellcome Trust Core Award Grant Number 090532/Z/09/Z, the John Fell Oxford University Press (OUP) Research Fund and the Li Ka Shing Foundation via a Oxford-Stanford Big Data in Human Health Seed Grant.

